# Genetics-informed bidirectional mapping of glycemic traits and brain phenotypes

**DOI:** 10.64898/2026.01.08.698365

**Authors:** Boxing Liu, Yunfan Zhang, Jianwei Chen, Chao Li

**Affiliations:** School of Science and Engineering, University of Dundee, UK; Department of Clinical Neurosciences, University of Cambridge, UK; Department of Applied Mathematics and Theoretical Physics, University of Cambridge, UK; School of Medicine, University of Dundee, UK

## Abstract

Dysglycaemia is linked to brain atrophy, white-matter disruption and dementia risk, yet the directionality and mechanisms of glycemic-brain coupling remain unclear. Here we integrate large-scale GWAS of fasting glucose, fasting insulin, 2-hour glucose, HbA1c and type 2 diabetes with multimodal brain imaging phenotypes and major brain disorders using bidirectional two-sample Mendelian randomization. Across 4,040 tests, we identify 54 glycaemia and brain causal associations (FDR P < 0.05) and uncover a timescale-dependent hierarchy: short-term glycemic traits map to distributed macrostructure and functional-network, whereas long-term glycemic burden preferentially implicates long-range association and commissural white-matter pathways with focal prefrontal-paracentral vulnerability. Reverse analyses reveal 44 brain IDPs and glycaemia effects. Multi-trait Bayesian colocalization and tissue/cell-type eQTL integration nominate shared causal loci enriched in glial and neurovascular pathways. Individual-level UK Biobank analyses validated glycaemia-associated brain structural alterations and non-linear risk patterns for incident outcomes. Together, our results provide a genetically anchored atlas of bidirectional glycemic-neural relationships, linking metabolic dysregulation to multiscale brain vulnerability and disorder risk.

## Introduction

Blood glucose regulation is essential to human physiology, and disruptions in glycemic control affect more than half a billion people worldwide. Beyond systemic metabolic and vascular consequences, chronic hyperglycemia and insulin resistance have been associated with cortical thinning, reduced white-matter integrity, cognitive decline, and elevated risk of stroke and dementia. Yet, the specific anatomical pathways and molecular processes through which glycemia relates to brain organization remain insufficiently defined.

Biologically, the relationship between glycemic physiology and the brain is bidirectional. Hypothalamic, limbic, and autonomic circuits regulate appetite, insulin secretion, hepatic glucose production, and energy expenditure, while circulating glucose influences neuronal excitability, glial metabolism, and long-term brain integrity through oxidative, inflammatory, and microvascular pathways. These reciprocal links provide biological plausibility for the systematic coupling of glycemic and brain traits. Accordingly, associations between glycemic traits and brain integrity detected in population imaging genetic studies likely reflect genetically patterned, system-level bidirectional relationships. However, simple association studies cannot determine the direction of these relationships and the shared genetic mechanisms. This underscores the need for genetically informed approaches that leverage inherited variation to reduce confounding and provide more directionally interpretable evidence than conventional association studies.

Large-scale resources such as the UK Biobank, BIG-KP, and ENIGMA now provide multimodal MRI phenotypes with sufficient depth to characterize brain organization at regional, microstructural, and network levels. MRI-derived phenotypes, spanning cortical morphology, subcortical structure, white-matter microstructure, and functional connectivity, are highly heritable and biologically meaningful endophenotypes that capture brain organization. At the same time, glycemic traits have well-characterized genetic architectures, with large-scale GWAS from MAGIC and DIAGRAM identifying loci involved in insulin secretion, glucose sensing, lipid metabolism, and pancreatic β-cell function. FinnGen provides harmonized genetic data and information on neuropsychiatric disorders from a genetically isolated population. Integrating these resources creates a unique opportunity to investigate whether directional links exist between glycemic variation and neural circuits.

To address these gaps, we integrate GWAS of glycemic traits, multimodal neuroimaging, and neuropsychiatric disorders within a genetically informed framework using bidirectional two-sample Mendelian randomization (MR). We apply multi-trait colocalization tests to determine whether glycemic and brain phenotypes share common genetic loci. Additionally, eQTL-based, cell- and tissue-resolved annotation localizes these signals to specific biological pathways and brain cells. Using individual-level UK Biobank data, we also evaluate the risks of cognitive and clinical outcomes associated with glycemic burden and brain architecture. Our study presents a comprehensive, genetically anchored atlas of glycemic–neural relationships, establishing an integrative foundation for understanding how glycemic variation aligns with brain architecture and contributes to the risk of neuropsychiatric disorders.

## Results

### Study overview

We hypothesized that physiological variation across the glycemic spectrum, captured by fasting glucose (FG), fasting insulin (FI), 2-hour glucose (2hGlu), hemoglobin A1c (HbA1c), and type 2 diabetes (T2D), demonstrates genetically informed directional causal associations with brain structure, function, and risk of brain disorders. As these glycemic traits reflect different physiological domains, where acute (2hGlu) and subacute (FG, FI) glycemic variations capture short-term fluctuation, while chronic exposure (HbA1c, T2D) reflects long-term burden, we further anticipated that they may map onto distinct or partially overlapping neural circuits. In parallel, we investigated whether variation in brain structure or functional networks exhibits genetically informed causal associations with glycemic traits.

To evaluate these hypotheses, we synthesized large-scale GWAS of glycemic traits, multimodal brain imaging–derived phenotypes (IDPs), and neuropsychiatric disorders within a bidirectional Mendelian randomization (MR) framework. This enabled genetically proxied, directionally interpretable associations between glycemic traits and brain phenotypes. Multi-trait Bayesian colocalization and cis-eQTL integration identified shared loci and potential regulatory mechanisms, while individual-level analyses in the UK Biobank validated phenotypic associations and tested for non-linear dose-response patterns. We also performed mediation analyses to validate specific brain structures in linking these relationships. Finally, we derived polygenic and imaging-informed predictors of brain disorders and tested their generalizability in external cohorts.

Collectively, this multimodal strategy enabled us to construct a fine-grained, genetically anchored map of glycemic–neural relationships across macrostructure, microstructure, functional connectivity, and clinical outcomes.

Quantitative glycemic traits (FG, FI, 2hGlu, HbA1c) were obtained from the MAGIC consortium (~281,416 individuals without diabetes), and T2D data were from DIAGRAM (69,033 individuals). Brain imaging phenotypes were derived from the UK Biobank, including 617 IDPs: 257 macrostructural measures (cortical and subcortical volumes) and 360 microstructural indices (white-matter tracts) from up to 33,224 participants. Functional connectivity (FC) IDPs were derived from a meta-GWAS of 47,276 individuals, encompassing 191 reproducible regions across Yeo 7 cortical and subcortical-cerebellar networks. GWAS summary data for 11 brain disorders (six neurological and five psychiatric) were sourced from the FinnGen project (sample sizes 9,725-161,405), covering stroke, Alzheimer’s and Parkinson’s diseases, migraine, bipolar disorder, and major depression disorder.

Multi-trait Bayesian colocalization (multi-coloc) identified loci with shared causal variants, integrating brain tissue and cell-type cis-eQTL data from GTEx (N ≤ 209), MetaBrain (N ≤ 8,613), and single-cell eQTL datasets (N = 373). Functional annotation incorporated partitioned heritability (LDSC), tissue- and cell-type enrichment, and developmental expression data (BrainSpan), cross-referenced with draggability resources.

### Causal effects of glycemic traits on brain structure and function

We identified 54 genetically informed causal associations (P_FDR_ < 0.05) across 4,040 MR tests, spanning 14 macrostructural, 12 microstructural, and 3 functional connectivity (FC) phenotypes. Across modalities, a coherent timescale-dependent pattern emerged: short-term glycemic traits (FG, FI, 2hGlu) were linked to distributed cortical and functional network variation, whereas chronic traits (HbA1c, T2D) were associated with diffuse, long-range white matter vulnerability and focal prefrontal–paracentral cortical effects. These patterns suggest complementary but distinct pathways through which short-term glycemic fluctuations and long-term burden manifest in the brain.

#### Macrostructure

Across cortical territories, higher glycemic levels showed a consistent directional association with reduced cortical thickness and smaller regional volumes, most prominently within the prefrontal–cingulate–paracentral circuit, which spans the FPN, SMN, and SAN/DMN interfaces. Both short-term and long-term glycemic traits converged on prefrontal regions. Short-term traits produced broader, multi-lobar associations, while chronic traits exhibited more focal genetic vulnerability, concentrated within prefrontal and paracentral hubs.

FG and 2hGlu demonstrated convergent associations across prefrontal, parietal, and limbic cortices. For example, both traits were linked to reduced thickness in the right pars triangularis (FG: IVW β = −0.19, 95% CI = −0.30 to −0.08, P < 0.001; 2hGlu: IVW β = −0.18, 95% CI = −0.26 to −0.10, P < 0.001) and right frontal pole (FG: IVW β = −0.19, 95% CI = −0.29 to −0.08, P < 0.001; 2hGlu: IVW β = −0.14, 95% CI = −0.22 to −0.06, P < 0.001). In addition, FG is related to the volume and thickness of the right superior frontal gyrus, the thickness of the right rostral middle frontal, and the volume of the right pars triangularis. Similarly, 2hGlu shows causal associations with the thickness of the right pars opercularis and the medial orbitofrontal cortex. Notably, FG is also associated with bilateral paracentral cortex, part of the SMN, while 2hGlu effects extended into the limbic system (rostral and caudal anterior cingulate), parts of the DMN and SAN, respectively. FI showed focal associations with the left inferior temporal gyrus and the right amygdala, as well as global cortical volume (MaskVol-to-eTIV ratio, IVW β = −0.39, 95% CI = −0.59 to −0.19, P < 0.001).

By contrast, HbA1c and T2D were characterized by more regional associations, primarily in prefrontal and paracentral cortices. The effect of HbA1c is more focused on the prefrontal lobe, i.e., right superior frontal, right rostral middle frontal, right inferior frontal (pars opercularis/triangularis) gyri, and right frontal pole. Similarly, T2D showed focal effects in the paracentral cortex (IVW β = −0.07, 95% CI = −0.11 to −0.03, P < 0.001).

Together, these findings suggest that short-term fluctuations drive broader cortical variation, while long-term exposure leads to targeted vulnerability in the prefrontal and paracentral regions, with global implications.

#### Microstructure

In general, higher glycemic trait levels were consistently associated with lower ICVF, FA, and MO, and higher MD and OD, which are diffusion biomarkers indicative of reduced white matter integrity associated with axonal injury, myelin loss, and increased extracellular water. Further similar timescale-dependent patterns were observed.

Chronic exposure (HbA1c, T2D) broadly associated with long-range white-matter backbones that support cerebral information transfer, including association (uncinate fasciculus, inferior and superior fronto-occipital fasciculus), commissural (corpus callosum body, forceps minor and tapetum), projection (anterior corona radiata), and brainstem (cerebral peduncle, medial lemniscus, superior and inferior cerebral peduncle) fibers. Of all the causal pairs, HbA1c shows the largest effect on the OD value of superior fronto-occipital fasciculus (IVW β = 0.25, 95% CI = 0.11 to 0.39, P < 0.001), recapitulated by T2D.

In contrast, short-term traits (FG, 2hGlu) produce spatially restricted effects, primarily within the superior and inferior cerebellar peduncles, the medial lemniscus, the tapetum, and the cingulum cingulate gyrus that link the cerebellar-SMN system with LIN and associative cortices. For instance, FG levels were associated with ISOVF of the cingulum cingulate gyrus (IVW β = −0.24, 95% CI = −0.35 to −0.12, P < 0.001).

These findings suggest a spatial gradient of glycemic impact, from focal, short-term disruptions in LIN and cerebellar tracts to widespread, chronic degeneration across long-range association and commissural fibers.

#### Functional connectivity

The forward MR revealed associations restricted to short-term traits (2hGlu and FG) with SMN and FPN, indicating that acute/subacute glycemic elevations selectively modulate intrinsic neural coupling. Specifically, higher FG strengthened connectivity between the subcortical-cerebellar and SMN-DAN networks in the cerebellum and the postcentral and precentral regions in both net 21 (IVW β = 0.21, 95% CI = [0.10 to 0.33], P < 0.001) and net 55 (IVW β = 0.20, 95% CI = [0.10 to 0.30], P < 0.001), while higher 2hGlu was associated with an increased amplitude of the CEN and DMN (IVW β = 0.18, 95% CI = [0.10 to 0.25], P < 0.001) in the temporal region. These findings, consistent across functional network parcellations (21 vs. 55 brain regions), suggest that glycemic spikes selectively enhance functional coupling in cerebello-thalamo-SMN circuits and upregulate CEN-DMN interactions (fig 3C).

**Fig. 1.**
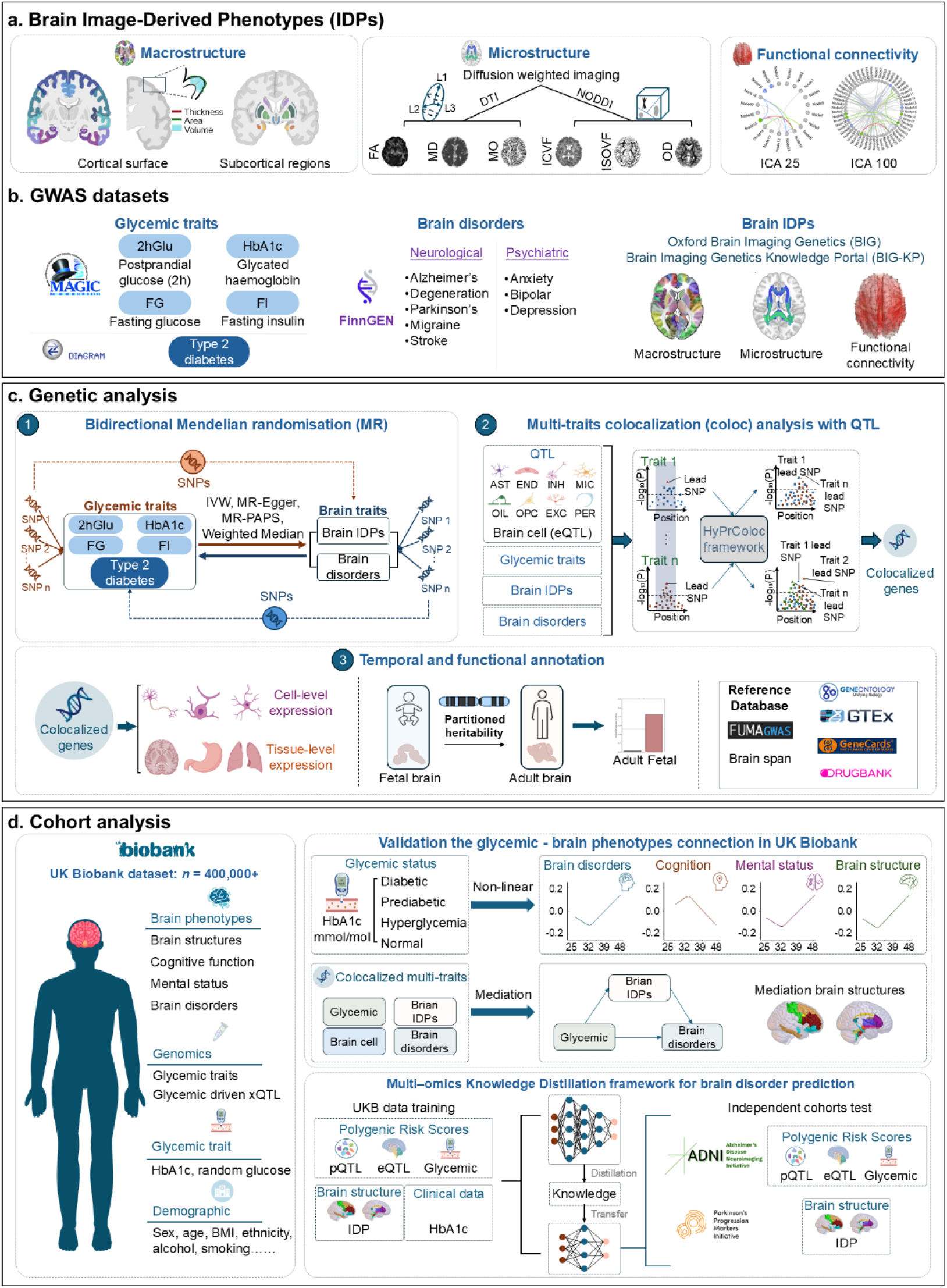
Overview of the study design and analysis.

**Fig. 2.**
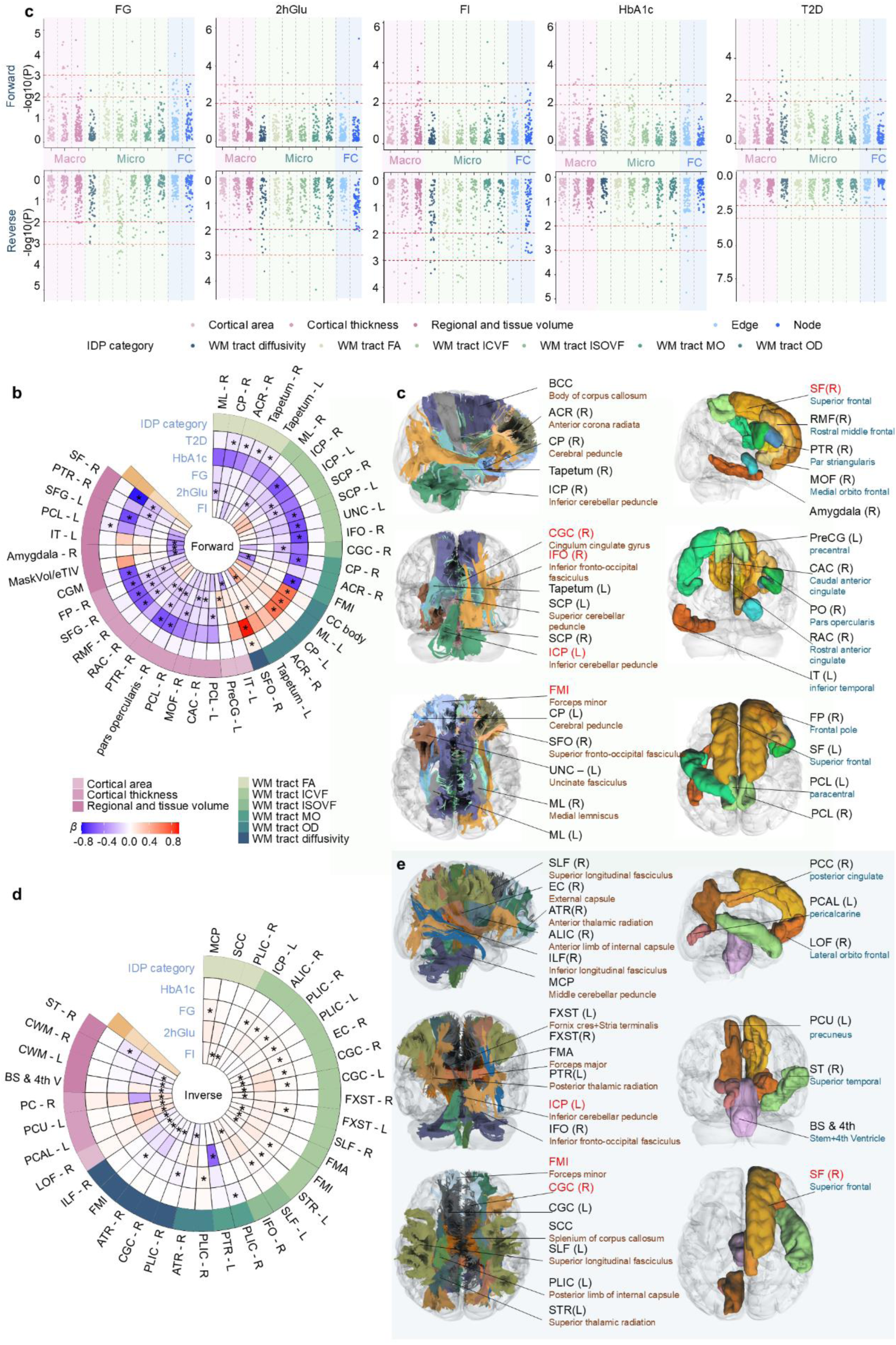
Phenotypic glycemic traits-brain structures causal associations. a. overview of the bidirectional mendelian randomization (MR) analysis across all imaging categories including forward and reverse MR. Macro: macrostructure; Micro: microstructure; FC: functional activity. b. significant result of forward MR (FDR P<0.05) c. details of IDP of forward MR. d. significant result of reverse MR (FDR P<0.05). e. details of IDP of reverse MR

**Fig. 3.**
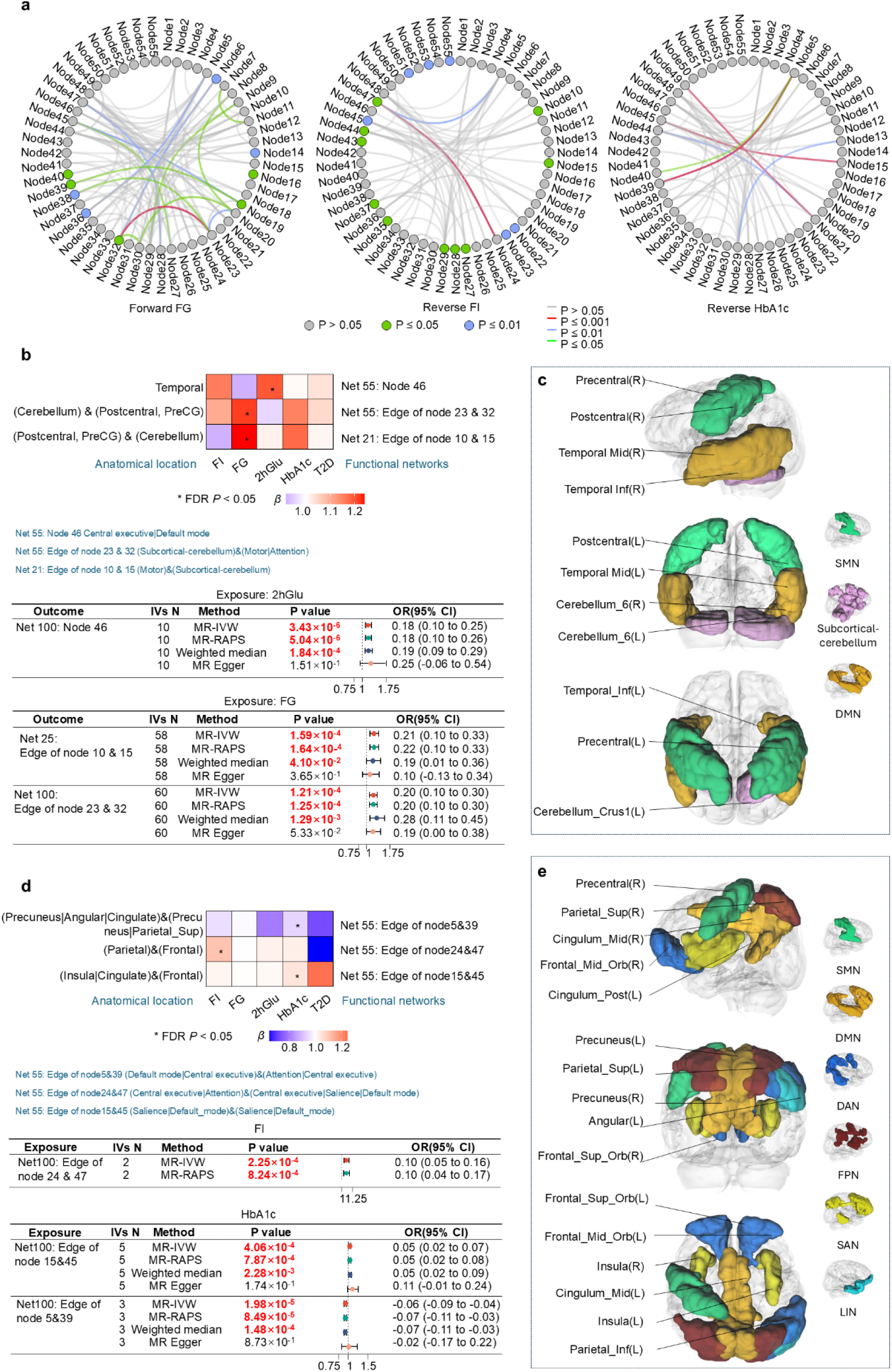
MR analysis between glycemic traits and functional connectivity

**Fig. 4.**
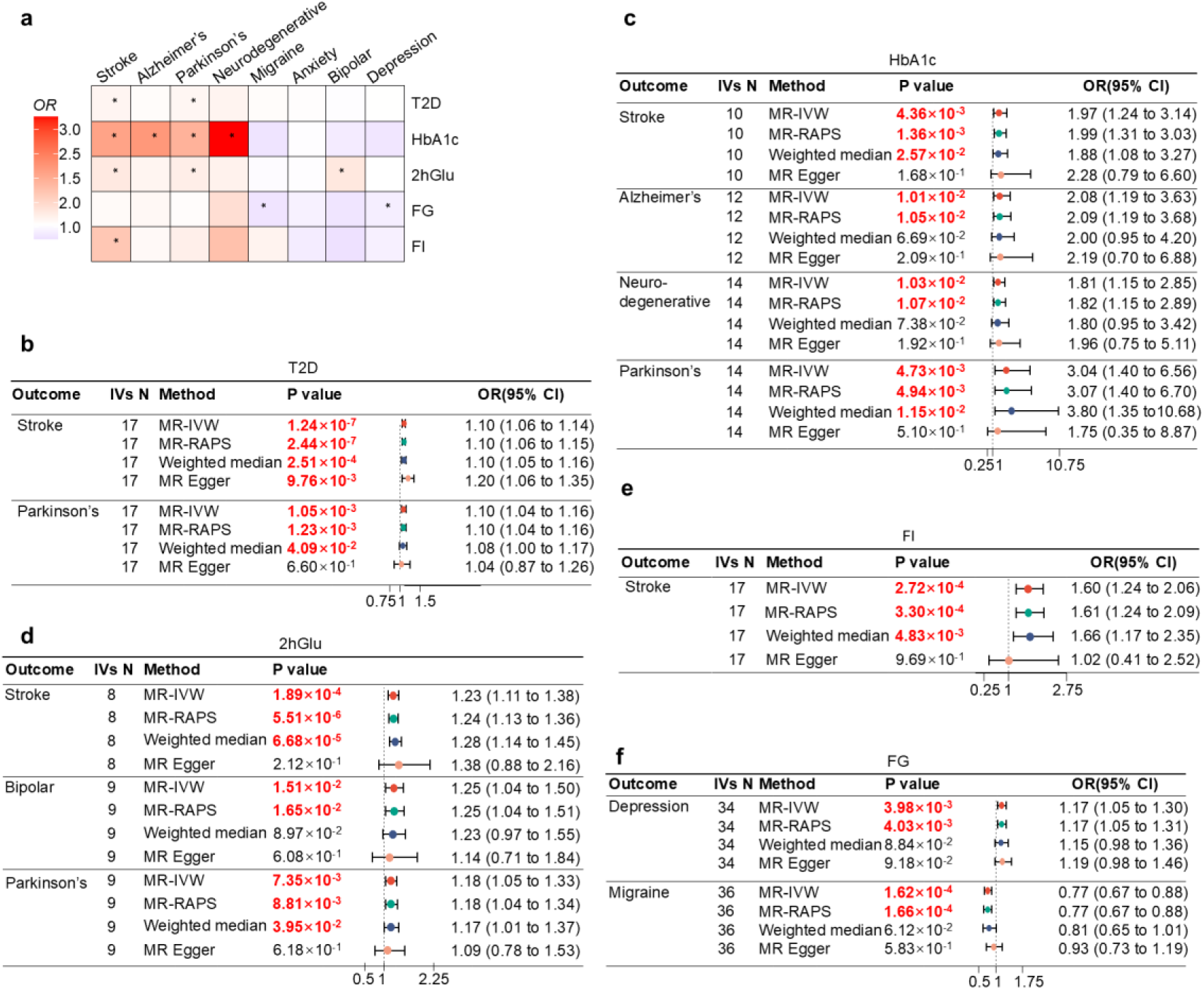
significant MR analysis between glycemic traits and brain disorder (FDR P<0.0.5)

**Fig. 5.**
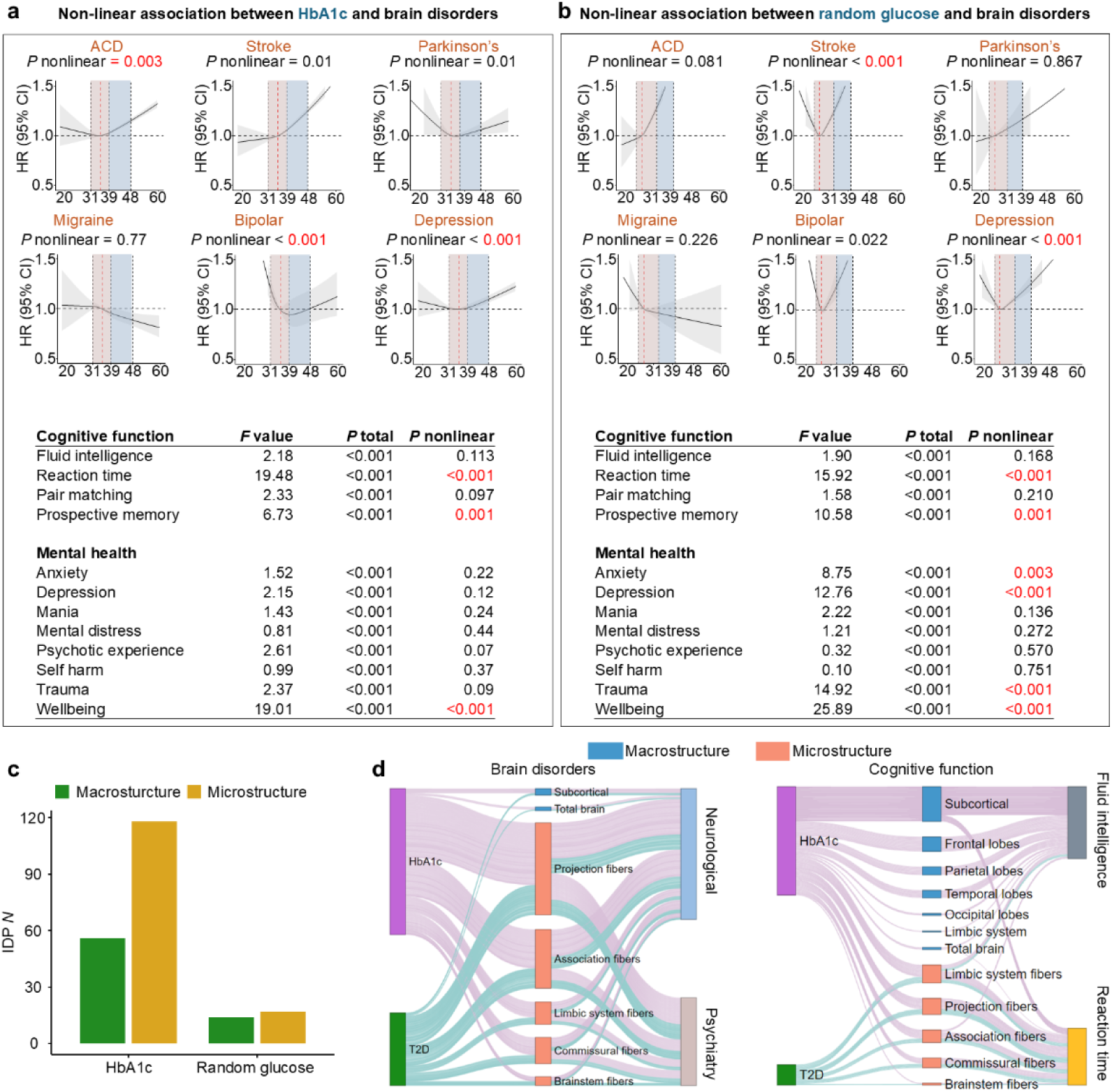
Cohort analysis in UKB. a. non-linear associations between HbA1c and brain healthy traits. b. non-linear associations between random glucose and brain healthy traits. c number of significant non-linear associations between random glucose and brain IDPs. d. mediation analysis between HbA1c, T2D and brain disorders and cognitive function.

Across structural and functional imaging, short-term glycemic traits are associated with broad cortical involvement and functional modulation, whereas chronic glycemic burden primarily links to long-range white matter tracts, reflecting the effects of distinct mechanisms. Nevertheless, their convergence on common neural targets, particularly the prefrontal cortex, suggests shared pathways that link glycemic profile with executive control, integrative processing, and higher-order metabolic regulation.

### Causal effects of brain structure and function on glycemic traits

The reverse MR identified 44 significant pairs of IDPs and glycemic traits, revealing distributed cortical, thalamic, and cerebellar pathways whose structural and functional variation is directionally linked to glycemic regulation. These associations included eight macrostructures, 18 microstructures, and three FCs (*P*_FDR_ < 0.05), which were more frequently observed for FI and FG (FI: 23/44 IDPs; FG: 16/44 IDPs; 2hGlu: 1/44 IDPs; HbA1c: 3/44 IDPs). Effects were most prominent in the prefrontal, cingulate, and parietal cortices, as well as in association tracts, including thalamocortical and cerebellar connections, implicating higher-order cognitive, sensorimotor, and autonomic circuits in shaping glycemic dynamics. Functionally, these structural substrates corresponded to FPN, SAN, and SMN networks, suggesting that variation in attention- and stress-regulatory circuits contributes to individual differences in glycemic physiology.

#### Macrostructure

Several cortical and subcortical regions showed directional associations with glycemic traits, particularly FI, including the right lateral orbitofrontal, right posterior cingulate, left pericalcarine, left precuneus, stem and 4th ventricle, and bilateral cerebellar white matter. These structures encompass the frontal lobe, limbic system, occipital lobe, and parietal lobe, as well as subcortical regions that overlap with FPN, LIN, DMN, Motor, and visual function networks, aligning with their known involvement in stress regulation, interoception, autonomic control, and sensory integration. In addition, a higher volume of the superior temporal was linked to lower FG, consistent with a role in auditory-sensory integration and homeostatic feedback.

A higher area of the superior frontal gyrus was associated with higher T2D risk, suggesting altered executive-attentional circuitry that may contribute to impaired metabolic regulation through stress, planning, or behavioural pathways.

#### Microstructure

White-matter integrity demonstrated robust associations with FI and FG, especially in the forceps minor, cingulum–cingulate gyrus, and anterior/posterior thalamic radiations, linking prefrontal and SMN cortices to thalamic relay nuclei. These findings suggest that thalamocortical connectivity, which is central to attention, arousal, and sensory gating, serves as a structural substrate influencing glycemic control.

Associations involving the inferior and middle cerebellar peduncles further support cerebellar contributions to autonomic modulation and anticipatory metabolic regulation, consistent with emerging evidence of cerebellar–visceromotor involvement in homeostasis.

#### Functional connectivity

Higher-order FC analyses revealed that stronger coupling between FPN, DAN, and SAN in parietal and frontal lobes was associated with higher FI (IVW β = 0.10, 95% CI = [0.05 to 0.16], P < 0.001) levels, suggesting with an arousal- and stress-related modulation (Figure 3 A). Similarly, enhanced SAN-DMN coupling (insula, cingulate, and frontal) was linked to higher HbA1c (IVW β = 0.05, 95% CI = [0.02 to 0.07], P < 0.001). In contrast, stronger DMN-FPN connectivity (precuneus, angular, cingulate and precuneus, parietal_sup) corresponded to lower HbA1c (IVW β = −0.06, 95% CI = [−0.09 to −0.04], P < 0.001). The SAN-DMN synchrony may reflect stress-related dysregulation, while adaptive functional coupling within executive and default mode circuits may support metabolic stability.

Together, these reverse causalities demonstrate that genetically influenced variation in structural and functional circuits regulating stress, autonomic output, and feeding behavior may contribute to differences in glycemic physiology. Our findings provide large-scale evidence that higher-order thalamocortical and cerebellar systems contribute to glucose regulation, extending classical models of hypothalamic-LIN control toward cortical-subcortical feedback mechanisms.

### Bidirectional causal effect between glycemic traits and brain disorders

We next examined bidirectional genetically informed associations between glycemic traits and 15 major brain-related disorders to determine whether the structural and functional patterns identified above translate into clinical outcomes, and whether brain disorders show directional associations with glycemic regulation.

Across the glycemic spectrum, chronic glycemic exposure (HbA1c, T2D) displayed the broadest and most consistent risk profile, associated with elevated neurovascular and neurodegenerative risks. In contrast, short-term glycemic exposure (FG, 2hGlu, FI) displayed a more heterogeneous pattern, with protective associations in several psychiatric conditions but risk elevations for vascular and neurodegenerative phenotypes. These results align with earlier findings, showing that chronic burden predominantly affects long-range white-matter, while short-term fluctuations align with functional modulation.

Specifically, higher HbA1c was causally associated with increased risk of stroke (IVW OR = 1.97, 95% CI = [1.24 to 3.14], P = 4.36×10^−3^), Alzheimer’s disease (AD: IVW OR = 2.08, 95% CI = [1.19 to 3.63], P = 0.01), Parkinson’s disease (PD: IVW OR = 3.04, 95% CI = [1.40 to 6.56], P = 4.73×10^−3^), a neurodegeneration composite outcome (IVW OR = 1.81, 95% CI = [1.15 to 2.85], P = 0.01), consistent with a pervasive neurovascular and neurodegenerative burden. T2D showed a similar pattern, with genetically proxied liability increasing the risk of stroke (IVW OR = 1.10, 95% CI = [1.06, 1.14], P < 0.001) and Parkinson’s disease (IVW OR = 1.10, 95% CI = [1.04, 1.16], P=1.05×10^−3^), aligning with basal ganglia and cerebrovascular pathways implicated in the structural analysis.

2hGlu aligned with HbA1c and T2D, shown causal effects on risks of stroke (IVW OR = 1.23, 95% CI = [1.11 to 1.38], P < 0.001), Parkinson’s disease (IVW OR = 1.18, 95% CI = [1.05 to 1.33], P = 7.35×10^−3^), and bipolar (IVW OR = 1.25, 95% CI = [1.04 to 1.50], P = 0.02), implying that postprandial dysglycemia may extend risk to affective and cognitive domains. Higher FI was only associated with increased stroke risk (IVW OR = 1.60, 95% CI = [1.24 to 2.06], P < 0.001), consistent with established links between insulin resistance, endothelial dysfunction, and vascular risk.

In contrast, FG levels displayed several protective associations with migraine (IVW OR = 0.77, 95% CI = [0.67 to 0.88], P < 0.001) and depression (IVW OR = 0.85, 95% CI = [0.77 to 0.95], 3.98×10^−3^), indicating that stable or lower short-term glycemic levels may influence psychiatric vulnerability differently from chronic exposure, paralleling the distinct patterns observed in functional networks.

The reverse MR identified that a higher risk of PD was associated with an increased risk of T2D and a higher level of 2hGlu. This suggests that Parkinson’s disease may increase genetic liability to T2D and 2hGlu, reflecting brainstem-basal ganglia contributions to autonomic and metabolic control observed in the structural and connectivity analyses.

### Integrated bidirectional interpretation

Collectively, these findings delineate a genetically anchored bidirectional relationship between glycemic traits and brain organisation. In the forward direction, glycemic traits show directional associations with brain integrity: short-term traits (FG, FI, 2hGlu) relate to widespread cortical and functional-network variation, whereas chronic glycemic burden (HbA1c, T2D) is linked primarily to long-range association and commissural white-matter pathways, reflecting distinct temporal sensitivities of neural systems to glycemic variation.

In parallel, the prefrontal-thalamic-cerebellar circuits exhibit directional associations with glycemic traits in the reverse analyses, aligning with their recognised roles in stress regulation, autonomic control, and behavioural modulation of energy balance. These convergent associations across cortical, subcortical, cerebellar, and white-matter systems support a multiscale pattern of glycemia–brain coupling mediated by inherited genetic variation.

Extending these patterns to clinical phenotypes, the results reveal a genetically informed relationship between glycemic traits and brain disorders: chronic glycemic burden aligns with increased risk of neurovascular and neurodegenerative diseases, whereas short-term glycemic fluctuations show stronger associations with psychiatric outcomes, mirroring their functional-network effects. The reverse MR further indicates that disorders involving autonomic, basal ganglia, or brainstem pathways exhibit directional associations with glycemic traits, reinforcing a broader bidirectional framework in which systemic glycemic profiles and distributed neural circuits mutually reflect lifelong, genetically mediated variation.

### Pleiotropy of genetic variants across glycemic traits, brain structures, and disorders

To clarify the mechanisms underlying MR-identified causal effects, we performed multi-trait Bayesian colocalization to determine whether glycemic traits and brain phenotypes share the same causal variant. Evidence of pairwise colocalization was defined as posterior probability (PP) > 0.7. In multiple loci, including the FADS1/FADS2 and ZBED3-AS1 regions, glycemic traits and white-matter IDPs colocalized with eQTLs from recent large-scale meta-analysis of brain tissue and cells, indicating that both traits arise from a common upstream genetic signal. This pattern supports vertical pleiotropy, consistent with a pathway in which genetic variation modulates glycemia and downstream glial processes that, in turn, shape white-matter organization. Thus, colocalization provides mechanistic coherence for the MR findings by linking causal effects of glycemia on brain structure to specific genes, tissues, and cellular pathways.

#### Glycemic traits and brain phenotypes

We identified 11 genomic regions showing pleiotropy between glycemic traits and brain structures. Notably, short-term glycemic traits are predominantly colocalized with macrostructures, whereas long-term traits are mainly colocalized with microstructures. Particularly, 2hGlu and FI are exclusively colocalized with macrostructures. The strongest evidence was observed at rs174559 (FADS1/FADS2), where HbA1c colocalized with the left uncinate fasciculus (PP=0.98) and the right inferior fronto-occipital fasciculus (PP=0.97), implicating lipid desaturation and myelination processes in glycemia-related white-matter integrity. At rs6878122 (ZBED3-AS1) and rs34228231 (NUP160), FG shared causal variants with the left inferior cerebellar peduncle (PP=0.96), indicating astrocytic WNT signaling (ZBED3-AS1) and nucleocytoplasmic transport mechanisms (NUP160) that may mediate short-term glycemic effects on cerebellar–limbic connectivity.

#### Glycemic traits and brain disorders

We identified nine genomic regions jointly influencing glycemic traits and brain disorders. Chronic traits (HbA1c, T2D, FI) predominantly colocalized with neurological diseases, while FG showed stronger links to psychiatric disorders. Specifically, the strongest evidence is that T2D shares causal variants with PD at rs356219 (SNCA) (PP = 0.99), a canonical susceptibility locus for PD. The second strongest evidence is that HbA1c and stroke colocalized at rs10757283 (ANRIL) (P = 0.99), indicating an epigenetic neurovascular mechanism (ANRIL-CDKN2A/B) linking chronic glycemic burden to cerebrovascular risk. Moreover, FI and stroke colocalized at rs7903146 (TCF7L2), indicating a shared causal variant that links β-cell function to cerebrovascular risk via a TCF7L2/Wnt regulatory axis. In contrast, FG colocalized with BD (PP=0.71) and MDD (PP=0.90) at rs174564 (FADS2), suggesting that lipid-myelin pathways may link metabolism and mood disorders.

#### Brain cell and tissue eQTLs

We systematically tested whether causal variants for glycemic traits overlap with eQTLs in eight different brain cell types. Across eight major types, 20 genomic regions showed colocalization. A glia-centric genetic architecture was evident, with strong enrichment in oligodendrocytes (5 regions) and oligodendrocyte precursor cells (OPCs)-committed oligodendrocyte precursors (COPs) (5 regions), followed by astrocytes (3 regions) and microglia (3 regions). Of note, OPCs/COPs were exclusive to long-term traits (HbA1, T2D, PP = 0.83-0.97). Neuronal signals were broadly linked to both short-term and long-term traits, specifically excitatory neurons with FG, FI, and T2D (four regions), and inhibitory neurons with FG and HbA1c (three regions). Endothelial enrichment (one region) was confined to HbA1c.

Moreover, at brain tissue level, we identified that FG colocalized with basal ganglia, cerebellum, cortex and hippocampus covering five genomic regions (PP = 0.80-0.97). Similarly, HbA1c colocalized with basal ganglia, cortex, hippocampus and spinal cord covering seven genomic regions (PP = 0.72-0.99). Specifically, the strongest evidence is that HbA1c colocalized with cortex (PP=0.99) at rs117890063 (LITAF), an endosomal/lysosomal membrane protein that regulates TNF-α–mediated inflammatory signalling and the endocytosis and degradation of receptors. We identified that T2D colocalized with cortex (PP=0.99) at rs11243150 (SSR1), which broadly expressed across most tissues, including the brain, and in neurons and glial cells it participates in the biosynthesis of diverse secreted and membrane proteins.

#### Peripheral tissues eQTLs

To further investigate whether causal variants for glycemic traits overlap with eQTLs in whole body tissue. The results revealed that HbA1c has the largest range of colocalized tissues (PP = 0.71-0.97), encompassing 29 tissues, including adipose, liver, skeletal muscle, pancreas, and whole blood. T2D colocalized with 18 tissues (0.70-0.99), also including adipose, liver, muscle skeletal, pancreas and whole blood, which are consistent with HbA1c. For FG, we identified 12 colocalized tissues (PP=0.73-0.97), including adipose.

### Cohort analysis

To validate MR-derived associations at the phenotypic level, we conducted individual-level analyses using UK Biobank longitudinal and imaging data.

We used a generalized linear model (GLM) to explore associations between HbA1c and brain structure, cognitive function, mental status, and brain disorders. HbA1c showed significant associations with all cognitive domains, except for pairs matching, after adjusting for age, sex, ethnicity, BMI, alcohol, smoking, Townsend index, education, and Haemoglobin. Specifically, higher levels of HbA1c were associated with decreased cognitive function, including reaction time, prospective memory, and fluid intelligence (P < 0.001). The strongest effect was observed for increasing reaction time (β = 0.02).

Then we investigated the association between HbA1c and mental status. We observed that higher levels of HbA1c will significantly decrease all mental health (anxiety, depression, psychotic experience, trauma, and wellbeing) except for mania, mental distress, and self-harm status after multi-test correction.

We further investigated the association between HbA1c and brain structures. The results indicated that a higher level of HbA1c was associated with brain atrophy and damaged brain connectivity. Specifically, higher levels of HbA1c will decrease the cortical thickness, volume, and area. Moreover, higher levels of HbA1c were associated with lower FA, MO, and ICVF, and higher levels of MD, OD, and ISOVF values in white matter tracts, resulting in consistent MR analysis.

#### Prospective association between HbA1c and brain disorders

Given the prospective design of the UK Biobank, we first examined the longitudinal associations between HbA1c and incident brain disorders. We first categorized the population into diabetic (HbA1c > 48 mmol/mol), prediabetic (39 ≤ HbA1c < 48 mmol/mol), normoglycemic (31 ≤ HbA1c < 39 mmol/mol), and hypoglycemic (HbA1c < 31 mmol/mol) groups. Multivariable Cox proportional hazards models were used to estimate hazard ratios (HRs) for incident outcomes, with the normoglycemic group as the reference. For major neurological outcomes, including all-cause dementia (ACD), Parkinson’s disease (PD), and stroke, participants in the abnormal HbA1c groups generally had higher risks compared with the normoglycemic group. For ACD, prediabetic significant increase the risk of ACD by 11% (HR = 1.11, 95% CI = 1.05 to 1.17, P < 0.001), diabetic significant increase the risk of ACD by 84% (HR = 1.84, 95% CI = 1.70 to 1.92, P < 0.001). In particular, the diabetic group has the highest risk of brain disorders. The results revealed that diabetic significant increase the risk of stroke by 79% (HR = 1.79, 95% CI = 1.67 to 1.92, P < 0.001) and PD by 68% (HR = 1.68, 95% CI = 1.49 to 1.89, P < 0.001).

For psychiatric outcomes, we observed a more complex pattern. Both hypoglycemic and, whereas the prediabetic group tended to have a lower risk of these disorders relative to normoglycemic participants. diabetic groups showed higher risks of depression and bipolar disorder. To further clarify the association within the non-diabetic range, we conducted additional logistic regression analyses restricted to individuals without diabetes. In this subset, higher HbA1c levels were significantly associated with a decreased risk of bipolar disorder (β=-0.11, *P*=0.006), supporting the notion that the relationship between HbA1c and psychiatric outcomes is unlikely to be strictly linear.

To formally assess potential non-linear dose–response relationships, we then modelled HbA1c as a continuous variable using restricted cubic splines (RCS) within the Cox Proportional Hazard model, with full adjustment for covariates. The tests for non-linearity were significant for several outcomes (all P for non-linearity < 0.001), indicating that linear models did not adequately capture the association. The spline curves revealed J-shaped associations of HbA1c with ACD and stroke, with the lowest risk observed around the normoglycemic to mildly elevated range and higher risks at both low and particularly high HbA1c levels. In contrast, HbA1c showed U-shaped associations with bipolar disorder and depression, characterized by elevated risks at both low and high HbA1c, and a risk around intermediate levels, consistent with the lower risk observed in the prediabetic group. For PD, the RCS analysis also suggested a U-shaped pattern (P for non-linearity = 0.01), with slightly increased risks at both ends of the HbA1c distribution and a more pronounced risk elevation at higher levels, aligning with the elevated PD risk in the diabetic group. Taken together, these findings suggest that the relationships between HbA1c and brain disorders are largely non-linear, exhibiting distinct J- and U-shaped patterns across neurodegenerative, cerebrovascular, and psychiatric outcomes.

#### The mediation of brain structures between glycemic traits and brain disorders

Previous MR analysis identified 42 brain structures with robust evidence of causal effects from at least one glycemic trait. We therefore considered these 42 phenotypes as glycemia-sensitive regions of interest (ROIs). Individual-level analyses were conducted using HbA1c and T2D, which were the only glycemic measures available in the cohort. Firstly, we focused on ROIs with MR evidence specifically for HbA1c (18 ROIs) or T2D (six ROIs). We conducted mediation pathway analyses for brain disorders using the structural equation model (SEM) to investigate the mediating role of brain structures. All results were conducted multi-test corrections.

Totally, 11 ROIs (11/18) demonstrated a mediating effect on the relationships between HbA1c and the risk of ACD (seven ROIs), bipolar (three ROIs), MDD (seven ROIs), stroke (five ROIs) and PD (two ROIs), while no brain structures mediated the link between HbA1c and migraine. It is noted that total grey matter volume mediated the association between HbA1c and all brain disorders. Besides total grey matter volume, two macrostructural ROIs (volume and thickness of superior frontal) and four microstructural ROIs (SFOF MD, UF ICVF, BCC OD and IFO ICVF) mediated the association between HbA1c and ACD. For stroke, four microstructural ROIs (SFOF MD, UF ICVF, BCC OD and CP OD) mediated the associations. SFOF MD mediated the association between HbA1c and PD. In contrast, three microstructural ROIs (SFOF MD, UF ICVF, IFO ICVF) and four macrostructural ROIs (pars opercularis, pars triangularis, rostal middle frontal and superior frontal) mediated the associations between HbA1c and MDD. Similarly, UF ICVF and IFO ICVF mediated association between HbA1c and bipolar.

For T2D, totally, 5 ROIs (5/6) demonstrated a mediating effect on the relationships between T2D and the risk of ACD (2 ROIs), bipolar (three ROIs), MDD (three ROIs), stroke (five ROIs), and PD (one ROI). Especially, all ROIs were microstructures.

Additional exploratory analyses evaluated associations between HbA1c, T2D and ROIs that had been prioritised by MR for other glycemic traits (fasting glucose, fasting insulin or 2-h glucose), acknowledging that these traits capture overlapping aspects of glycemic regulation.

## Discussion

In this study, we integrated large-scale GWAS of glycemic traits with multimodal brain phenotypes and clinical outcomes to construct a bidirectional map of glycemic–neural coupling. Using harmonized summary statistics and individual-level validation, we identified a timescale-dependent hierarchy: short-term traits (FG, FI, 2hGlu) associate with widespread cortical and functional changes, whereas chronic burden (HbA1c, T2D) impacts long-range white matter. Reverse analyses revealed thalamocortical and cerebellar circuits influencing glycemic regulation beyond hypothalamic-limbic pathways. Disease-level MR supported these directions, showing chronic burden linked to neurovascular and neurodegenerative risk, and short-term traits with mixed psychiatric diseases. Colocalization highlighted glial and neurovascular mechanisms, particularly oligodendrocyte-lineage lipid metabolism (FADS1/FADS2) and astrocytic pathways (ZBED3-AS1) as molecular mediators of metabolic-neural coupling.

Our results reinforce prior MR evidence that hyperglycemia and T2D increase stroke and neurodegenerative risk, linking higher HbA1c and fasting insulin to cognitive decline and brain aging. The observed alterations in the DMN-CEN/SAN, alongside disrupted visual and temporal connectivity, mirror population-level findings of cortical thinning and white-matter loss associated with chronic hyperglycemia. Further, our results align with epidemiological links between depression risk and metabolism, indicating that glycemic imbalance disrupts both cognitive and affective circuits. Compared to existing evidence, this study advances the field through: (i) fine-grained multimodal mapping of macrostructure, microstructure, and functional coupling; (ii) bidirectional inference uncovering metabolic-neural feedback via thalamocortical and cerebellar circuits; and (iii) mechanistic colocalization identifying glial and metabolic pathways (FADS1/FADS2, ST6GAL1, G6PC2) linking systemic glycemia to brain organization. Collectively, these results define a systems-level framework in which short-term fluctuations shape functional coupling, and chronic glycemic load drives structural compromise through glial lipid and metabolism pathways.

We acknowledge inconsistencies in the prior research, such as mixed associations between T2D and Alzheimer’s disease and occasional protective effects of fasting glucose. These differences likely stem from variations in study design, trait specificity, and cohort characteristics. Nonetheless, hyperglycemia and T2D consistently increase stroke risk and relate to reduced white-matter integrity and brain volume, aligning with evidence that insulin resistance impairs cognition. Our fine-grained mapping, which focuses on prefrontal-sensorimotor regions and thalamocortical tracts, diverges from studies that emphasize occipital or hippocampal areas, likely due to improvements in imaging resolution, the inclusion of both micro- and macro-structural features, and bidirectional modeling. Furthermore, prior work has highlighted reduced DMN coherence; we extend this to reveal cerebello-thalamo-sensorimotor and executive-salience coupling, the first genome-informed FC analysis linking neural feedback to systemic glycemia beyond affective domains.

The convergence on prefrontal and sensorimotor systems suggests that executive control, arousal/stress gating, and motor-autonomic integration share neural substrates for metabolic and cognitive control. White-matter vulnerability under chronic glycemia highlights myelin/lipid pathways as potential interventional targets, while short-term network modulation suggests that acute cerebrovascular-metabolic coupling drives psychiatric phenotypes. Clinically, the atlas of effects supports: (i) risk stratification using glycemic and imaging markers; (ii) mechanism-guided trials (e.g., testing whether glycemic optimization, insulin sensitizers, or lipid–myelin modulators attenuate fronto–callosal and thalamic-tract alterations); and (iii) cross-disciplinary prevention, integrating mental health interventions to stabilize glycemic dynamics.

Our study has limitations. MR relies on key assumptions (relevance, independence, exclusion restriction); despite the use of pleiotropy-robust estimators and sensitivity analyses, horizontal pleiotropy cannot be fully ruled out. Instrument strength varies across traits (notably for some FC IDPs), potentially reducing power. The sample ancestry was predominantly European, which limited the generalizability of the results. IDPs and FC parcellations, while reproducible, remain operational definitions of brain organization; between-cohort harmonization and measurement error may attenuate effects. Disease phenotypes pooled from GWAS cohorts can include heterogeneity in case definition and comorbidity. Validation in UK Biobank mitigates some concerns but is observational and subject to residual confounding; mediation analyses are supportive but not definitive of individual-level pathways. Finally, our approach is largely cross-sectional in genetic time and cannot pinpoint disease duration or exposure timing without explicit longitudinal designs.

Future directions should (i) integrate continuous glucose monitoring (CGM) with longitudinal MRI to resolve temporal coupling between glycemic dynamics and network activity; (ii) expand diversity (ancestry, age, sex) to assess portability and modifiers; (iii) build multimodal predictors that combine polygenic scores, glycemic indices, and IDPs for early identification of neurovascular and neurodegenerative risk; (iv) evaluate interventional targets via drug-gene MR and trials (e.g., GLP-1R agonists, SGLT2 inhibitors, remyelination strategies) with imaging endpoints; (v) pursue mechanistic studies linking oligodendrocyte lipid metabolism, astrocytic glucose transport, and mitochondrial/ER stress to tract-specific vulnerability.

In summary, our results support a unified, bidirectional model in which short-term glycemic fluctuations modulate cortical structure and functional coupling, whereas chronic glycemic burden preferentially compromises long-range white matter integrity. The anatomical, genetic, and cell-type specificity of these effects provides a coherent substrate for mechanism-informed prevention and therapeutics at the interface of metabolic and brain health.

## Methods

### Ethics declarations

This research was approved by the ethics committee of XXX. We used the brain-imaging data under UK Biobank application 52802, with XXX. as principal investigator. The UK Biobank received ethical approval from the North West Multi-Centre Research Ethics Committee. Other publicly available GWAS data have their own ethical approvals; please see the respective references.

### Data source and study population

#### UK Biobank data

Our study was based on data from the UKB, a large-scale longitudinal cohort including over 500,000 participants aged 37–73 years from 2006 to 2010 ^1^. All participants provided electronic signed consent for their data to be used in health-related research. In this study, after excluding participants with lumped event dates (UKB data-coding 819) and brain disorders (N = 85,996), 478,377 participants remained in the analysis. For brain disorders, we included first occurrence, summary diagnosis and cancer diagnosis, influenced brain health.

#### GWAS data of glycemic traits

Here, we tested the performance of bidirectional MR on GWAS datasets of glycemic-related traits. Firstly, we obtained GWAS summary statistics of HbA1c, FI, FG and 2hGlu from a meta-analysis of previous GWAS from the meta-analyses of glucose and insulin-related traits (MAGIC) consortium^2^ with a total sample size of 281,416 individuals of European ancestry. We also included summary-level GWAS data of T2D derived from diabetes genetics replication and meta-analysis (DIAGRAM)^3^, which contain 34,840 cases and 114,981 controls.

#### GWAS data of brain IDPs

Brain IDPs. We used the GWAS summary statistics of IDPs processed by Smith et al.^4^ with a sample size of almost 40,000 individuals of European ancestry from the UK Biobank. Based on reliable segmentation and meaningful measurement ^5–8^, 617 brain structural IDPs were selected from the 3,913 original releases, including 214 IDPs in the cerebral cortex, 43 IDPs in subcortical regions and 360 IDPs of white matter integrity. We further classified these IDPs into 12 regional categories and 9 measures ^6,8^. The tools or analyses used to get these measurements are reliable and are described. In addition, we obtained 191 functional IDPs derived from rfMRI, including 76 nodes and 115 edges ^9^.

1. Cortical measurements were extracted by the FreeSurfer tool based on the Desikan–Killiany atlas ^7^.
2. Subcortical measurements were extracted by the FreeSurfer tool based on the automatic subcortical segmentation ^6^.
3. The features of white matter connections were estimated from the diffusion magnetic resonance imaging based on two complementary analyses, tract-based spatial statistics ^8^ and probabilistic tractography ^10^. Details of these neuroimaging measures and processing methods60 are provided by the online reference of UK Biobank (https://biobank.ctsu.ox.ac.uk/crystal/crystal/docs/brain_mri.pdf). All GWAS summary statistics on IDPs were obtained from the Oxford Brain Imaging Genetics (BIG40) web browser (https://open.oxcin.ox.ac.uk/ukbiobank/big40/).
4. Functional brain regions and corresponding functional connectivity were characterized via spatial independent-component analysis (ICA)^11,12^. Two parcellations with different dimensionalities ^9,13^ (25 and 100 regions, respectively) was separately applied in spatial ICA, and we focused on the 76 (21 and 55, respectively) regions that had been previously confirmed to be nonratification. All GWAS summary level data derived from Brain Imaging Genetics Knowledge Portal (BIG-KP) web browser (https://bigkp.org) ^9^.

#### GWAS data of diseases

We collected nine neuropsychiatric disorders with publicly available GWAS summary statistics, including neurological disorders: stroke, Alzheimer’s diseases, Parkinson’s diseases, Neurodegenerative disorder, dementia and migraine and psychiatric disorders: anxiety, bipolar and depression. All of this data were derived from FinnGEN (www.finngen.fi/en/access_results), which is the specific genetic makeup of the Finnish population. In the GWAS of selected, well-studied diseases, we were able to identify several new associations with a fraction of the cases compared with the largest published GWAS. The sample sizes for these nine disorders ranged from 500,000 individuals.

#### eQTL data

In this study, we performed cell-type–resolved colocalization analyses by integrating single-cell brain eQTL data. Summary-level single-cell eQTL results were obtained from Bryois et al.^14^, which, to our knowledge, is currently the only resource reporting eQTLs across all major cell types in the adult human brain using single-cell RNA sequencing. Briefly, this dataset provides eQTL summary statistics for eight cell types—astrocytes, endothelial cells, excitatory neurons, inhibitory neurons, microglia, oligodendrocytes, oligodendrocyte precursor cells and pericytes—derived primarily from the prefrontal and temporal cortices of 192 individuals of European ancestry. In the original study, expression levels for ~14,595 genes were quantified and ~5.3 million SNPs were analysed.

To replicate colocalization findings from the single-cell eQTL analyses and to further prioritise putative causal genes, we additionally leveraged bulk-tissue eQTL summary statistics from GTEx (v8)^15^. We focused on GTEx brain tissues sampled from regions comparable to those in the single-cell dataset, with broadly similar sample sizes. We also analysed selected peripheral tissues to assess whether colocalization between glycaemic traits and brain disorders extends beyond the brain. Across GTEx tissues, approximately 10.4 million SNPs were tested for association with gene expression in each tissue.

### MR analysis

#### Two-sample MR

We conducted two-sample MR^16^ analyses to estimate the causal effects of genetically predicted glycaemic traits on brain imaging-derived phenotypes (IDPs) and brain disorders using the TwoSampleMR package in R^17^. GWAS summary statistics for glycaemic traits were obtained from large-scale studies of individuals of European ancestry. For each glycaemic trait, we selected independent genetic instruments as SNPs associated at genome-wide significance (P < 5 × 10^−8^) and clumped for linkage disequilibrium (LD) at r^2^ < 0.001 within a 10,000 kb window using European reference panels from the 1000 Genomes Project. Inverse-variance weighted (IVW) MR under a multiplicative random-effects model was used as the primary estimator^18^. Because IVW estimates can be biased in the presence of horizontal pleiotropy, we applied four complementary MR approaches to assess robustness. MR-RAPS accounts for both systematic and idiosyncratic pleiotropy and is well suited to settings with many weak instruments ^19^. MR-Egger estimates the causal effect from the slope of the Egger regression and can provide consistent inference under the InSIDE assumption, even when all instruments exhibit pleiotropic effects^20^. The weighted median estimator yields consistent estimates provided that at least 50% of the weight comes from valid instruments^21^. The weighted mode estimator is consistent when the largest cluster of instruments (by weight) represents valid variants and can show reduced bias and type I error under relaxed instrument validity assumptions^22^. When only a single genetic instrument was available, we used the Wald ratio^23^. All analyses were implemented using the TwoSampleMR R package (v0.6.29)^17^ via the functions mr_ivw, mr_raps, mr_weighted_median, mr_egger_regression, mr_weighted_mode and mr_wald_ratio.

#### Bidirectional MR

We extended the above MR analysis to a bidirectional causal inference between glycemic traits and brain phenotypes. Forward MR analyses were performed with glycemic traits as exposures and brain phenotypes as outcomes. Conversely, reverse MR analyses were performed with brain phenotypes as exposures and glycemic traits as outcomes. We performed MR analyses in accordance with the STROBE-MR checklist^24^ and Burgess et al.’s guidelines^25^.

### Multi-trait colocalization analyses

As the instruments used in the current setting were identified based on their statistical associations with the protein level, we conducted another sensitivity analysis-colocalization, to investigate whether the genetic associations with both protein and phenotypes shared the same causal variants. We conducted colocalization analysis for each of the six proteins associated with one or more of the stroke outcomes to investigate whether the protein level and stroke outcome genetic associations are due to the same causal variants. We estimated the posterior probability (PP) of multiple traits sharing the same causal SNP simultaneously using a multi-trait colocalization (HyPrColoc) method8. Variants within a ±1 Mb window around the eQTLs with the smallest P value, with imputation (INFO)-score ≥0.8 and minor allele frequency (MAF) ≥0.01, were included. All variants across each of the datasets were harmonised to the same effect alleles prior to colocalization analyses. We conducted the colocalization analysis using the ‘HyPrColoc’ R package^26^.

